# Machine learning for single cell genomics data analysis

**DOI:** 10.1101/2021.02.04.429763

**Authors:** Félix Raimundo, Laetitia Papaxanthos, Céline Vallot, Jean-Philippe Vert

**Affiliations:** Google Research, Brain team, Paris, France; CNRS UMR3244, Institut Curie, PSL University, Paris, France; Translational Research Department, Institut Curie, PSL University, Paris, France; MINES ParisTech, PSL University, CBIO-Center for computational biology, Paris, France

**Author notes:** Equal contribution (Félix Raimundo, Laetitia Papaxanthos).

## Abstract

Single-cell omics technologies produce large quantities of data describing the genomic, transcriptomic or epigenomic profiles of many individual cells in parallel. In order to infer biological knowledge and develop predictive models from these data, machine learning (ML)-based model are increasingly used due to their flexibility, scalability, and impressive success in other fields. In recent years, we have seen a surge of new ML-based method development for low-dimensional representations of single-cell omics data, batch normalization, cell type classification, trajectory inference, gene regulatory network inference or multimodal data integration. To help readers navigate this fast-moving literature, we survey in this review recent advances in ML approaches developed to analyze single-cell omics data, focusing mainly on peer-reviewed publications published in the last two years (2019-2020).

## Introduction

With single-cell omics technologies getting wide-spread adoption, computational methods are urgently needed to process the large amounts of data they produce [1]. Machine learning (ML) approaches have recently demonstrated their fantastic potential to automatically process and learn from large amounts of high-dimensional data in fields such as computer vision or natural language processing [2]. They are therefore seen by many as a promising way to infer biological knowledge and develop predictive models from single-cell omics data, which provide high-dimensional characterization of large quantities of cells. Not surprisingly, the development of ML approaches to analyze single-cell omics data has been a very active field of research recently.

In this review we survey recent advances in ML approaches developed to analyze single-cell transcriptomic and epigenomic data, focusing mainly on peer-reviewed publications published in the last two years (2019-2020). This period witnessed active developments of new methods, in particular based on deep learning, to automatically extract information from large sets of single-cell data, tackling important problems such as batch normalization, multimodal data integration, automatic cell type classification, trajectory inference or gene network reconstruction. It is also a period where systematic benchmarks started to highlight the practical challenges associated to these methods, as well as their potential. With this review we hope to give the reader enough entry points to that fast-moving literature in order to grasp the current state-of-the-art and join its future developments.

### From raw data to useful representations

Raw single-cell transcriptomic count data, as well as their epigenomic counterparts, provide a high-dimensional and noisy description of each cell by assessing the activity of thousands of genes or DNA loci simultaneously. Transforming raw count data to a lower-dimensional representation of each cell using dimension reduction (DR) technique is a useful step to remove technical noise and prepare data for visualization, classification or further analysis tasks (Figure 1).

**Figure 1:**
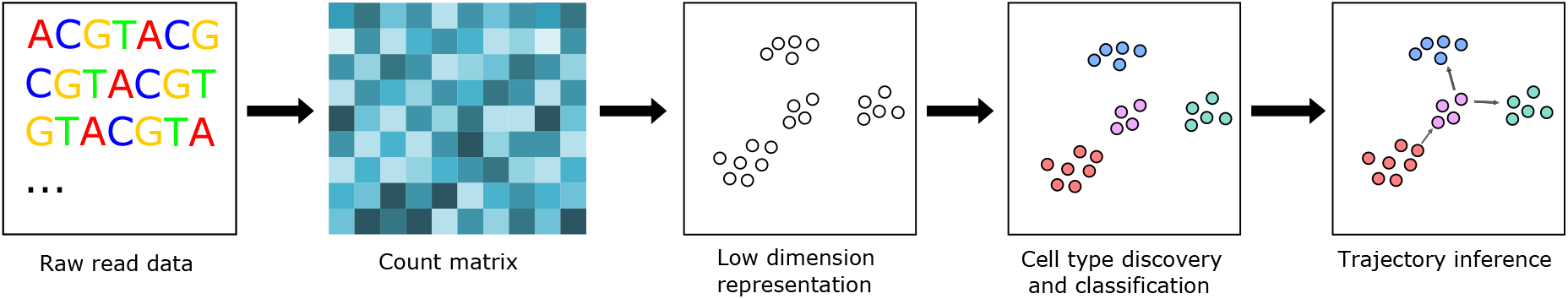
Standard analysis pipelines using a single modality of single-cell omics data start by turning the raw sequencing reads into a matrix of cells×feature counts. This matrix is then used for dimension reduction, representing each cell by a vector of lower dimension (embedding). The embedding is then used as starting point for subsequent tasks such as visualization, cell type discovery, or trajectory inference.

While early and widely-used methods such as scran [3] and Seurat v2 [4] use standard principal component analysis (PCA) on log-transformed count data for DR, many new DR models have been proposed specifically for scRNA-seq data recently. A common theme has been to replace the implicit Gaussian noise assumption of PCA by explicit statistical models of raw count data, modelling for example overdispersion and zero-inflation due to dropout in the matrix factorization-based model ZinbWave [5], or heavy-tailed count distribution in the nonparametric Bayesian model of [6]. Several groups have also investigated the potential of (variational) autoencoders ((V)AE), a very popular class of deep learning-based DR models. In short, a (V)AE learns a low-dimensional representation of input data (cell transcriptomes in our case) that is sufficient to reconstruct the input data, using flexible neural network models to go from the input to the compressed representation (encoding), and from the representation to the input data (decoding). Several (V)AE models for scRNA-seq data have been proposed recently, include scVI [7^•^], DCA [8], SAVER [9] and scVAE [10]. Methods using hyperbolic geometry have also recently been developed [11^•^] (J. Ding et al. bioRxiv doi: https://doi.org/10.1101/853457). These models differ from each other by some modelling assumptions, such as the statistical model for count data in the decoder, or the prior distribution of the low-dimensional representation, but otherwise follow a similar architecture. An interesting property of these models is their computational scalability, as they are typically implemented with deep learning libraries designed to train models with millions or more input points.

Have deep learning-based (V)AE definitively imposed themselves as the best DR approach for scRNA-seq data? The answer is not so simple. Besides requiring large number of cells to learn parameters, (V)AE performance was shown to be very sensitive to arbitrary parameter choices [12], and [13] highlighted that with datasets of a few hundreds or thousands cells simpler models remain competitive and easier to use. The practical difficulty to correctly train complex ML models is not specific to (V)AE: another example is the “art of training” the popular t-distributed stochastic neighbour embedding (tSNE) model for visualizing scRNA-seq in two dimensions [14], that requires specific initialization and choices of hyperparameters. Once correctly trained, tSNE reaches the same performance as uniform manifold approximation and projection (UMAP), a model proposed to improve tSNE mapping of scRNA-seq [15, 14]. This highlights, again, both the potential and the difficulty to train some modern ML-based models, and raises in particular important concerns about making sure that all published results are reproducible and not overfitted to a given experiment.

Several DR methods for single-cell epigenomic data have also been proposed recently, either based on standard PCA models [16, 17], matrix factorization with latent Dirichlet allocation [18], or a VAE [19]. A recent benchmark highlights the importance of preprocessing, in particular how reads are binned into regions of interest and counted, for the success of these methods [20^•^].

One interesting idea to use complex models on small datasets is to leverage larger, already annotated, datasets to learn the embedding, using techniques from the field of transfer learning or domain adaptation. Embeddings learned by PCA and non-negative matrix factorisation (NMF) on datasets such as the Human Cell Atlas (HCA) have successfully been used in both scATAC-seq [21] and scRNA-seq [22, 23] on new unseen datasets and cell types, as well as used for denoising the new dataset [24]. Similarly the embeddings learned by (demoising) AEs on one dataset, have been shown to be useful on other datasets, both for clustering [25, 26^•^, 27, 28] and surface protein prediction [29]. One limitation of these methods is that the embedding is only learned on a single dataset, and applied to another dataset, without analyzing both in parallel. This limits the ability to train models on multiple datasets and thus truly leverage the mass of experiments in databases such as HCA.

The result of the DR is often fed to standard clustering algorithms, as reviewed in [30], in order to identify cell types, with these algorithms also being extremely sensitive to hyperparameter choices [31]. Once the cells are clustered, differential expression tools, benchmarked in [32], can be used to identify *de novo* marker genes.

The cells can also be matched to known cell types either by querying a reference database with tools such as Cell BLAST [33], scMap [34], scQuery [35] or CellFishing.jl [36] or by using standard supervised learning techniques as benchmarked in [37]. However these methods can be sensitive to batch effects, whose corrections are the subject of the following section.

### Batch correction and integration of heterogeneous scRNA-seq data

Instead of analyzing data of a single experiment, much can be gained by jointly analyzing single-cell transcriptomic data of many experiments, potentially coming from different labs, using different technologies, and following different experimental protocols. ML models are likely to benefit from analyzing more cells, but the risk of capturing batch effects and other confounding factors instead of biological knowledge is large and considered one of the grand challenges of scRNA-seq data analysis [1]. A number of models have been proposed to specifically perform jointly DR on heterogeneous scRNA-seq data, build a global graph or construct a common gene expression matrix, aiming to capture biology and ignore confounding effects (see Figure 2 and [38] for a comprehensive benchmark).

**Figure 2:**
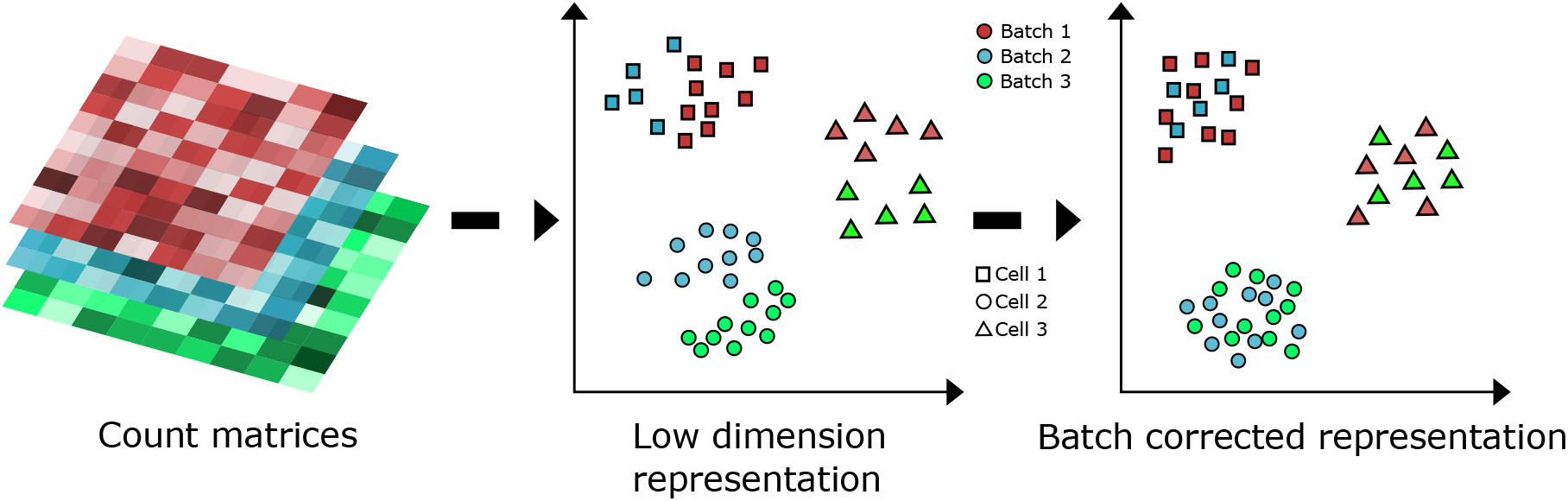
Different experiments of a similar modality (e.g., scRNA-seq) containing different number of cells can be integrated into a single unified view. At first, cells of the same type are separated by their batch, but after correction are perfectly merged together.

A first group of models learn a low-dimensional representation over a common space that is invariant to technical confounders. Among those, SAUCIE [39^•^] and scDGN [40] are deep-learning based, SAUCIE is an AE trained with a specific regularisation penalty on the latent codes to remove batch effects, and scDGN is a supervised adversarial neural network model trained to accurately classify cell types and discriminate against batches. scMC [41], Harmony [42] and SMNN [43] rely on a linear transformation to a lower dimensional space, clustering (shared nearest neighbour scheme, soft k-means or supervised mutual nearest neighbours) and post-processing of the low dimensional embeddings to both account for cell-cell similarities and remove batch-specific variations. Other models have an objective to build a joint graph connecting all measured cells, such as scPopCorn [44] which relies on PageRank and graph-k partitioning, and Conos [45] which exploits cell-cell similarity matrices and mutual nearest neighbours. These graph-based models allow for tasks such as cell annotation and information propagation along the network. However, the methods previously described hinder interpretability as they do not enable studying differentially expressed genes leveraging the multiple datasets. A third group of models attempt to tackle this problem by correcting for batch effects on the original count data. Among them, scAlign [46] uses paired AEs with a common latent space that conserves the cell-cell distances estimated in the count data, while BERMUDA [47] instead uses a regularisation penalty on cell clusters from different batches in the latent space, and scGen [48] combines VAEs and latent space vector arithmetics. scVI [7^•^] and trVAE [49] are so-called *conditional* VAE approaches that condition the decoder on an auxiliary batch variable to correct the data in the latent space. Based on variants of nearest neighbour search, scMerge [50] combines mutual nearest clusters and RUV-III factor analysis to remove unwanted factors from the count data, and Scanorama [51^••^] and Seurat v3 [52] rely on linear projection to a low-dimensional space and an efficient (mutual) nearest neighbour search to obtain matched cells in low-dimensional space that are used to build translation vectors in the high-dimensional space.

All methods cited above offer batch correction for scRNA-seq data, while scMC has also been proposed for scATAC-seq integration and SAUCIE for single-cell CyTOF measurements. While most methods need shared cell types across datasets to build anchor cells, SAUCIE, scPopCorn and scMerge can be used without. Finally, almost half of the methods are able to scale to datasets containing hundreds of thousands of cells.

### Learning trajectories, dynamics and regulation

Besides capturing the cellular heterogeneity of tissues and identifying cell types, single-cell omics data offers the possibility to learn about *dynamical* processes that shape this heterogeneity, such as cell cycle, differentiation, proliferation or tumorigenesis. From a data analytical point of view, this raises the question of inferring a dynamical model or at least the cellular trajectories from a snapshot of cells scattered at different time points along the dynamics. Since the first algorithms such as Monocle [53] were published in 2014 to infer trajectories and order cells using the notion of pseudotime, dozens of methods have been proposed. Recently proposed methods include GrandPrix [54], an efficient implementation of the Gaussian process latent variable model (GPLVM) to estimate pseudotimes and their uncertainty; STREAM [55], which estimates a low-dimensional set of curves, called the principal graph, to describe the cells’ pseudotime, trajectories and branching points; PAGA [56], a graph-based method to compute a graph representation of a set of cells, allowing visualization and dynamical interpretation at different resolutions; TinGa [57], which builds a graph to fit the single-cell omics data as well as possible using the Growing Neural Graph (GNG) algorithm; or Monocle 3 [58], the latest version of Monocle with new features such as learning trajectories with loops or point of convergence and better scalability. To help users choose a particular method for a given problem, [59^•^] published an impressive benchmark of trajectory inference methods, comparing 45 published algorithms on 110 real and 229 synthetic datasets. While no clear winner emerges in all situations, the benchmark is useful to understand the strengths and weaknesses of different methods in different settings.

A related problem is to infer the relationships between populations of cells captured at different time points along a dynamic process, such as developmental processes after induced pluripotent stem cell reprogramming observed through scRNA-seq profiles captured at half-day intervals [60^••^]. In that paper the authors develop a method, called Waddington-OT, to relate the populations of cells at different time points using the concepts and tools of optimal transport (OT), a mathematically well-established and fast-growing field in ML [61], particularly well adapted to compare populations of cells and model their evolution. With ImageAEOT, [62] show how OT combined with an autoencoder allows to predict the lineages of cells using time-labeled single-cell images.

While trajectory inference implicitly allows us to predict the future evolution of cells, some algorithms have also been proposed to explicitly infer the *velocity* of each individual cell’s transcriptomic profile. Following the pioneering work of [63], [64] proposed scVelo, a likelihood-based dynamical model for velocity inference from the ratio of spliced and unspliced mRNA. [65] propose another kernel-based velocity estimator, and show how gene regulatory networks (GRN) can be automatically inferred, although with modest accuracy, by training a sparse regression model to predict the velocity from gene expression levels. Another recent attempt to reconstruct GRN and more general gene networks from scRNA-seq data with an ML approach is the convolutional neural network for coexpression (CNNC) approach of [66], who represent each gene pair as a scatter plot of their expression levels across cells and train a standard CNN for 2D images on the resulting plots to learn pairwise relationships.

### Multimodal data integration

An important problem in single-cell omics data analysis is to integrate several modalities together, in order to enhance the performance of downstream tasks such as cell type labelling, identification of sub-populations, visualisation or regulatory network inference, as reviewed in [67, 68]. Several ML approaches have been developed for that purpose, for instance by characterizing cells across measurements, projecting multiple measurements into a common latent space or learning the missing modalities. Transcriptomics is typically one of the modalities that is integrated, together with chromatin accessibility [69, 52, 70], DNA [71], DNA methylation [72, 52], proteomic data [73, 74, 69, 75, 76] or CRISPR perturbations [77^•^, 78].

A first category of models assume that the correspondences between cells are known across modalities, with direct applications to co-assay data (Figure 3). Such methods learn a joint representation of each cell or a cell-cell similarity matrix that is used for downstream analyses by exploiting variants of VAEs such as totalVI (A. Gayoso et al., bioRxiv doi: 10.1101/2020.05.08.083337) and scMVAE [70], matrix factorisation-based models such as scAI [76] and MOFA+ [79], or k-nearest neighbour prediction to learn cell-specific modality weights as Seurat v4 (Y. Hao et al., bioRxiv doi: 10.1101/2020.10.12.335331). A second category of models do not require co-assays within individual cells and can be applied to independent multi-omics datasets originating from different cells. Current deep learning-based methods either rely on a pair of VAEs whose latent spaces are coupled through a specific penalty (K. D. Yang et al., arxiv.org/abs/1902.03515), or on learning low-dimensional representations minimising a tSNE loss for each view, coupled through a learned matching matrix (UnionCOM [74]). Other methods rely on NMF, to learn a low-dimensional space composed of specific and common factors (LIGER [75]), or cluster representatives of subpopulations of cells (DC3 [80^•^]). MMD-MA [73, 81] learns a joint latent representation where different modalities have a similar distribution using the theory of kernel methods. SCOT [69] uses OT to learn a joint distribution between cells from both views. clonealign [71] models the association between copy number features and gene expression leveraging mean field variational Bayes inference. While these methods can in theory be applied to any bi-modal omics dataset, hyperparameter selection is difficult when no co-assay data is available for MMD-MA, SCOT and UnionCOM. Among models that do not require co-assay data, some use weak supervision such as SCIM [72], an adversarial AE model that assumes that the cell types are known for a fraction of the cells and Seurat v3 [52], a canonical correlation analysis (CCA)-based model that relies on building anchor cells using mutual nearest neighbours. Applied to single-cell CRISPR screenings, scMAGeCK [78] relies on statistical analyses and MUSIC [77^•^] on topic modeling in order to link gene perturbations to cell phenotype. Finally, it is worth mentioning that some models require features to have a one-to-one correspondence between views [71, 52, 75, 77^•^, 78], which may not be the case systematically.

**Figure 3:**
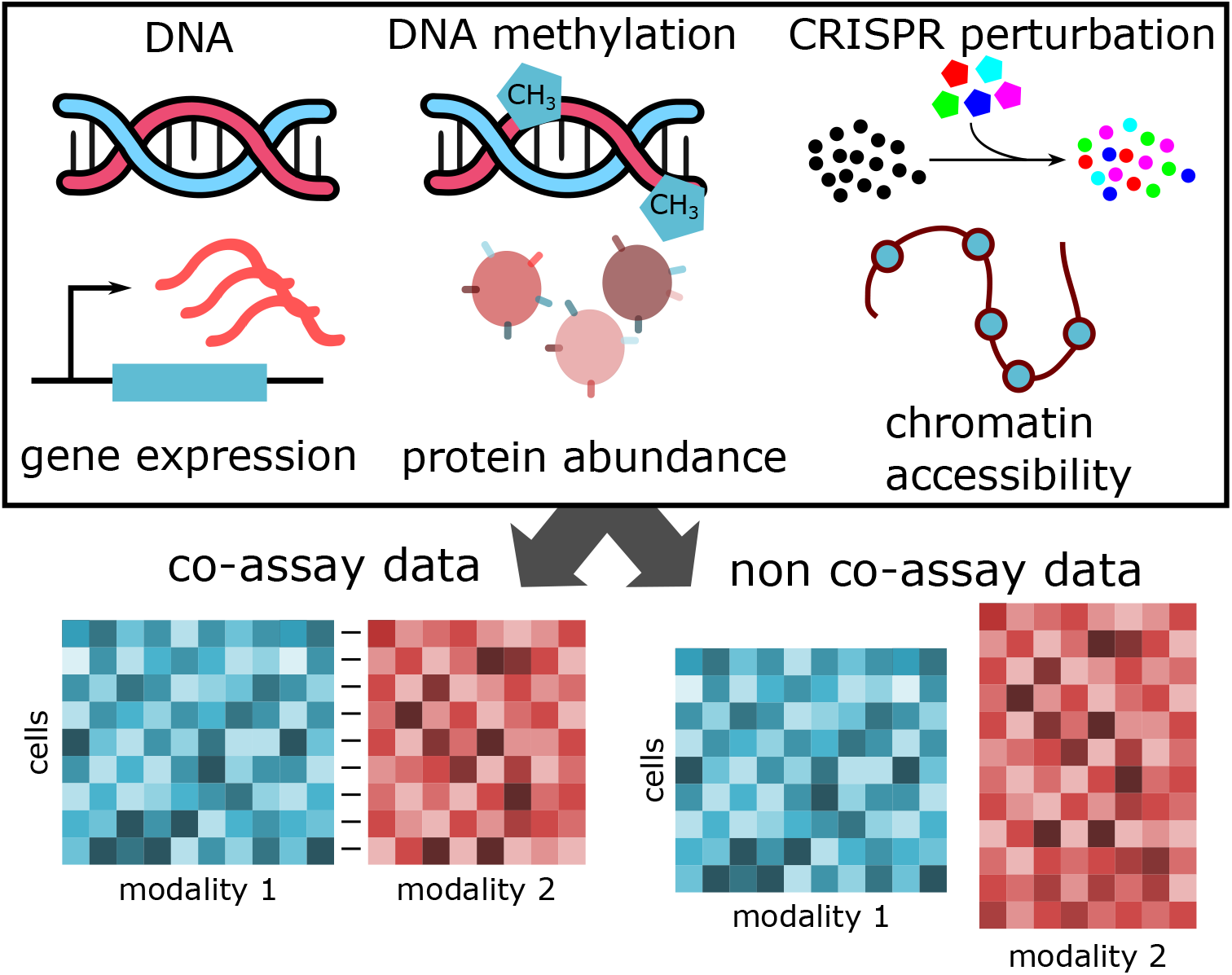
Single-cell modalities can take various forms, such as DNA, DNA methylation, CRISPR perturbations, transcriptomics, proteomics or chromatin accessibility. ML models developed for single-cell multimodal data integration assume that the correspondences between cells are either known (co-assay data) or not (non co-assay data) across modalities. In the case of non co-assay data, additional supervision signal might be used, such as cell types, correspondences between features or anchor cells.

While the diversity of models is large, most of them rely on finding a joint low-dimensional space that can be later used on downstream tasks. Most models combine two modalities and a few enable the integration of more than two, such as UnionCOM, MOFA+ and DC3, the latter also incorporating scHiC or bulk HiChIP datasets. Finally, the scalability of the models evolve conjointly with single-cell technologies, nowadays being able to handle tens or hundreds of thousands of cells [70, 72].

## Conclusion

Researchers are facing an exponential growth of approaches to deal with single-cell genomics data, with over 800 tools (scrna-tools.org) published for scRNA-seq analysis so far, many of which being based on ML approaches. A vast majority of ML-based tools have been straightforwardly imported from other fields, with some features unsuited for genomic challenges and to the reality of biological data - thereby not maximising their performance. In particular, a number of parameters, which have a strong impact on performance, need extensive training to be properly tuned, which is often unrealistic in the case of genomic data. It also raises questions of reproducibility that the scientific community should address, defining for example the processed datasets and variables that should be shared, i.e., random seed values or reduced dimensional spaces, in addition to the raw data. Whether ML models will in the near future make up for the current technical limitations of single cell genomics approaches - e.g dropouts, batch effects - remains uncertain. If current single-cell omics achieve genome-wide characterization of the transcriptome or epigenomes for example, these methods do not yet achieve single-locus/single-cell resolution due to the dropouts within datasets, leaving room for experimental and computational optimisation.

